# Characterization of the Doublesex/MAB-3 transcription factor DMD-9 in *Caenorhabditis elegans*

**DOI:** 10.1101/2022.09.30.510307

**Authors:** Rasoul Godini, Roger Pocock

## Abstract

DMD-9 is a *Caenorhabditis elegans* Doublesex/MAB-3 Domain transcription factor of unknown function. Single-cell transcriptomics revealed that *dmd-9* is highly expressed in specific head sensory neurons, with lower levels detected in non-neuronal tissues (uterine cells and spermatheca). Here, we characterized endogenous *dmd-9* expression and function in hermaphrodites and males to identify potential sexually dimorphic roles. In addition, we dissected the *trans-* and *cis*-regulatory mechanisms that control DMD-9 expression in neurons. Our results show that of the 22 DMD-9-expressing neuronal reporters we analyzed, only the neuropeptide-encoding *flp-19* gene is cell-autonomously regulated by DMD-9. Further, we did not identify defects in behaviors mediated by DMD-9 expressing neurons in *dmd-9* mutants. We found that *dmd-9* expression in neurons is regulated by four neuronal fate regulatory TFs: ETS-5, EGL-13, CHE-1, and TTX-1. In conclusion, our study characterized the DMD-9 expression pattern and regulatory logic for its control. We found that, as with other DMD TFs, DMD-9 likely acts redundantly to control neuronal development and function.

## Introduction

*C. elegans* is a microscopic nematode that generates two sexes: hermaphrodites and males. Despite core similarities, each sex possesses sexual-dimorphic morphology, nervous systems, and behavior (Lee and Portman 2007; Oren-Suissa *et al*. 2016; Cook *et al*. 2019). Hermaphrodites and males have 302 and 385 neurons, respectively, of which 294 neurons are common between sexes (Hobert 2005). Neuronal connectivity between some neurons is sex-specific and regulated following maturation (Oren-Suissa *et al*. 2016; Cook *et al*. 2019). In *C. elegans*, gender-specific characteristics are regulated by the sex determination pathway and members of the Doublesex/MAB-3 Domain (DMD)/DMRT family of Transcription Factors (TFs) (Shen and Hodgkin 1988; Yi *et al*. 2000; Wolff and Zarkower 2008; Oren-Suissa *et al*. 2016).

There are eleven DMD TFs encoded in *C. elegans* that regulate various sexual-dimorphic characteristics (Matson and Zarkower 2012). DMD TFs are expressed in both sexes but may exhibit sex-specific expression patterns. For example, MAB-3 is expressed in the male ADF thermosensory neurons, but not in hermaphrodites (Fagan *et al*. 2018). Likewise, DMD-3 is only expressed in the male PHC sensory neuron (Serrano-Saiz *et al*. 2017) and the DMD-5/DMD-11 TFs are only expressed in the male AVG interneuron (Oren-Suissa *et al*. 2016). Conversely, DMD-4 is expressed in the PHA and PHB neurons of adult hermaphrodites, but not adult males (Bayer *et al*. 2020). These sexual-dimorphic expression patterns regulate sex-specific characteristics and behaviors.

DMD TFs can regulate broad or specific sexual-dimorphic characteristics. MAB-3 is required for male V ray development, yolk development in males, male-specific cell lineages, and male behavior (Shen and Hodgkin 1988; Yi *et al*. 2000). MAB-23 is also required for the development of male sexual characteristics, including muscles, ray dopaminergic neuron patterning, axon pathfinding, and mating behavior (Lints and Emmons 2002). DMD-3 controls male-specific PHC neuron characteristics, such as development, synaptogenesis, and gene expression (Serrano-Saiz *et al*. 2017). DMD-4 controls synaptic formation in the PHA and AVG neurons in hermaphrodites (Bayer *et al*. 2020). DMD-5 and DMD-11 regulate AVG synaptic patterning and AVG-mediated male mating behavior (Oren-Suissa *et al*. 2016). DMDs may also regulate a behavior by impacting neuronal function. For example, DMD-10 regulates ASH-mediated responses to nose touch and high osmolarity (Durbeck *et al*. 2021).

Here, we focus on an uncharacterized member of the DMD TF family: DMD-9. Single-cell transcriptome analysis revealed that *dmd-9* is expressed in a subset of head sensory neurons (Taylor *et al*. 2021). DMD-9 is highly expressed in the BAG neurons, which sense oxygen (O_2_) and carbon dioxide (CO_2_) gases, regulate exploration and egg-laying behavior (Ringstad and Horvitz 2008; Zimmer *et al*. 2009; Juozaityte *et al*. 2017). DMD-9 is also expressed in AFD thermosensory neurons (Cassata *et al*. 2000), and AWB/ASE/AWC chemosensory neurons (Bargmann and Horvitz 1991; Bargmann *et al*. 1993; Troemel *et al*. 1997). Here, we studied DMD-9 expression throughout development in both sexes through endogenous tagging with GFP. We also investigated the potential functions of DMD-9 in neuronal fate determination and function in hermaphrodites and males. We also studied the *trans-* and *cis-*regulatory mechanisms that regulate *dmd-9* expression in the head neurons. We show that DMD-9 is not expressed in a sexually dimorphic manner in head neurons. However, it is expressed in non-neuronal sex-specific tissues; uterine cells in hermaphrodites and spermatheca in males. We found that expression of reporters in DMD-9-expressing neurons are not dependent on DMD-9, with the exception of the FMRF-amide neuropeptide *flp-19* in the BAG neurons. Finally, we identified *cis-* regulatory elements and fate determining TFs (ETS-5, EGL-13, CHE-1 and TTX-1) that drive *dmd-9* expression in head neurons. Taken together, our study characterized the DMD-9 expression pattern and regulation in both *C. elegans* sexes and found that, as with other DMD TFs, DMD-9 likely controls neuronal development and function in a redundant manner.

## Material and Methods

### Strains and maintenance of the *C. elegans* strains

All strains were maintained on nematode growth medium (NGM) plates at 20°C as described previously (Brenner 1974). Strains used in this study are listed in the Table S1. Animals were synchronized to specific ages by picking.

### Molecular analysis

#### CRISPR-Cas9

A coding sequence GFP-AID-TEV-FLAG was knocked-in to the last exon of *dmd-9* gene using CRISPR-Cas-9 (Dokshin *et al*. 2018). We designed the crRNA to guide the CRISPR-Cas-9 enzyme and a donor DNA fragment containing the insertion and overhangs to be inserted in the location of interest (https://sg.idtdna.com/sgRNA) (Table S2). CRISPR-Cas-9 mixture was injected into wild-type N2 animals and the F1 progeny were screened by genotyping and microscopy for GFP expression.

#### Cloning

Cloning and mutagenesis were performed using In-Fusion restriction-free cloning (Takara). Reporter gene constructs were cloned by inserting PCR amplified promoter elements into the pPD95.75 vector (Fire Vector Kit). Constructs were confirmed by Sanger sequencing.

#### Transgenic lines

Transgenic lines were generated by injecting constructs into young adult hermaphrodite N2 animals. *Pmyo-2::RFP*; *Punc-122::RFP* or *Punc-122::GFP* were used as co-injection markers. For reporter constructs, 30-50 ng/*μ*l of the construct was injected. For rescue lines, 1-5 ng/*μ*l of *ets-5* and *egl-13* fosmids was injected into *ets-5(tm866); him-8(e1489)* and *egl-13(ku194); him-8(e1489)*, respectively.

#### Auxin Inducible Degron (AID) analysis

Experiments were performed as described (Zhang *et al*. 2015). For the depletion of AID tagged proteins, we used 0.1mM or 1mM Auxin. For recovery experiments, we exposed the animals to 0.1mM auxin for 24h followed by 24h incubation on standard NGM plates. In all experiments, NGM plates were seeded with *Escherichia coli* OP50 bacterial lawns.

### Microscopy

Animals were mounted in sodium azide (NaN_3_) 25 μM. A Zeiss AXIO Imager M2 fitted with an Axiocam 506 mono camera and Zen 2 pro software (Zeiss) was used to take fluorescent and DIC images. Images were analyzed using Zen 2 pro software (Zeiss) and ImageJ v1.53c.

### Behavioral experiments

#### Chemotaxis

Chemotaxis behavior was performed on 90mm agar plates according to (Bargmann *et al*. 1993) with some modifications. For each replicate, 50 to 100 animals were used. In the case of animals carrying *him-8(e1489)* or *him-5(e1490)* mutations, 200-300 L4 males were transferred to a new plate to grow to adult. Depending on the chemical, different concentrations from 1:10 to 1:10000 (V/V) were freshly diluted in absolute ethanol (Merck). For NaCl chemotaxis, 1.0 and 0.25M concentrations were used (Bargmann and Horvitz 1991). To calculate the chemotaxis index the following formula was used:

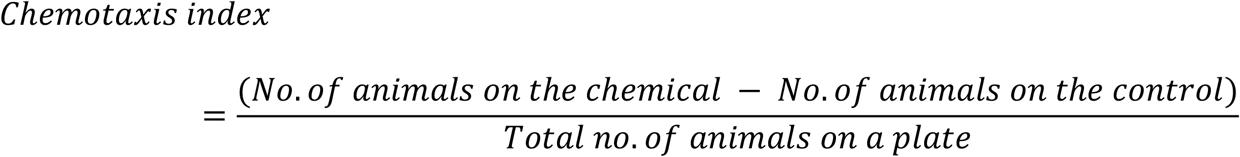

#### Gas sensing

Gas sensing was performed as previously described (Romanos *et al*. 2015). Animals were exposed to low concentrations of O_2_ (10% compared to 21% in control) or high concentrations of CO_2_ (1% compared to 0% in control) for 6 minutes. The proportion of gases was balanced by changing the N_2_ level. Animal motility was recorded with a 4-megapixel CCD camera (Jai) and analyzed using MatLab-based image processing to obtain instantaneous speed during continuous forward movements (Ramot *et al*. 2008; Tsunozaki *et al*. 2008).

#### Exploration

Exploration behavior was assayed as previously described (Flavell *et al*. 2013) with some modifications. Assay plates were 60 mm NGM plates uniformly seeded with *E. coli* OP50. Strains were grown well-fed for two generations at 20°C prior to each experiment. Exploration was assayed for young adult hermaphrodites and males for 16h and 2-3h, respectively. Animals were scored based on the count of squares entered in an 86-grid area.

#### Food leaving

Food leaving behavior was performed as previously described (Lipton *et al*. 2004). Experiments were performed on 90 mm plates spotted with 18 μL of *E. coli* OP50 (OD600 = 1.0) culture grown for 12-16h at room temperature. To synchronize animals, 20 to 30 L4 hermaphrodite or male animals of each strain were sex-segregated and matured for 12h. Individual animals were placed on the *E. coli* OP50 spot and scored for leaving events at 4, 8, 12, and 24h. A leaving event was scored if the animal reached 3 cm from the edge of the seeded bacteria (Lipton *et al*. 2004). To analyse the results, a survival model was applied to compare the ratio of leaver animals between different strains. Statistical significance of the experiment was calculated with the Gehan-Breslow-Wilcoxon test.

#### Egg-laying

Egg-laying behavior was performed as previously described (Ringstad and Horvitz 2008). Five adult hermaphrodites, synchronized from L4 for 30h, were transferred to assay plates containing 40 μL of freshly-coated *E. coli* OP50 culture seeded at the center of NGM plates and left to lay eggs for 1h at 20°C. The developmental stages of eggs laids were divided to stage 1 to 6 (Ringstad and Horvitz 2008) and analysed using a high-power dissecting microscope (Olympus SZX16). Distribution of egg developmental stages were analysed using Wilcoxon Mann-Whitney rank-sum test.

### Bioinformatic analysis

Transcriptomic analysis of *dmd-9* gene expression was obtained from (cengen.shinyapps.io/CengenApp/) (Taylor *et al*. 2021) and visualized using R programming. The JASPAR program (jaspar.genereg.net/ was used to predict TFs binding elements within the *dmd-9* promoter (Castro-Mondragon *et al*. 2022).

### Statistical analysis

Statistical analysis and visualization of plots were performed using GraphPad Prism® software version 8.0.2 (GraphPad Software Inc.). A *P*-value less than 0.05 was considered significant.

## Results

### DMD-9 is expressed in specific head sensory neurons in *C. elegans*

Single cell transcriptome data from L4 hermaphrodites shows that *dmd-9*, which encodes a member of the Doublesex/MAB-3 Domain (DMD) family of TFs, is expressed in neuronal and non-neuronal cells. High *dmd-9* expression was detected in the BAG, AFD, AWC^on^, AWC^off^, ASEL, and AWB neurons, with lower expression in ASER, and the ASGs (Fig. 1A-B) (Taylor *et al*. 2021). Additionally, *dmd-9* expression was detected in non-neuronal tissues, including uterine cells, vulva muscle, and sperm (Fig. 1A). Some members of the *C. elegans* DMD TF family (*dmd-3, dmd-5* and *dmd-11*) show sexually dimorphic expression patterns in *C. elegans* (Oren-Suissa *et al*. 2016; Serrano-Saiz *et al*. 2017). To investigate potential sex-specific expression pattern of DMD-9, we used CRISPR-Cas9 to endogenously tag the DMD-9 C-terminus with GFP and studied expression throughout development in both hermaphrodites and males (Fig. 1C-E). We detected DMD-9::GFP expression in 10 head neurons of hermaphrodites at the L4 stage (Fig. 1D). We also observed weak and inconsistent expression in other head neurons and transient weak expression in the tail of hermaphrodites and males at the L4 stage (not analyzed). No overt differences were observed in DMD-9::GFP expression in neurons of hermaphrodites and males (Fig. 1D). We also detected DMD-9::GFP expression in non-neuronal tissues. Transient DMD-9::GFP expression was observed in the uterine cells at specific sub-stages of the L4 stage. DMD-9 is not expressed from L4.0 to L4.3, with expression first detected in the early L4.4 stage, with the highest level during L4.4 before gradual loss by the L4.9 stage (Mok *et al*. 2015)(Fig. 1E). DMD-9::GFP was not detected in adult uterine cells (Fig. 1E). We also detected DMD-9::GFP expression in the spermatheca of hermaphrodites and males (Fig. 1E).

**Fig 1.**
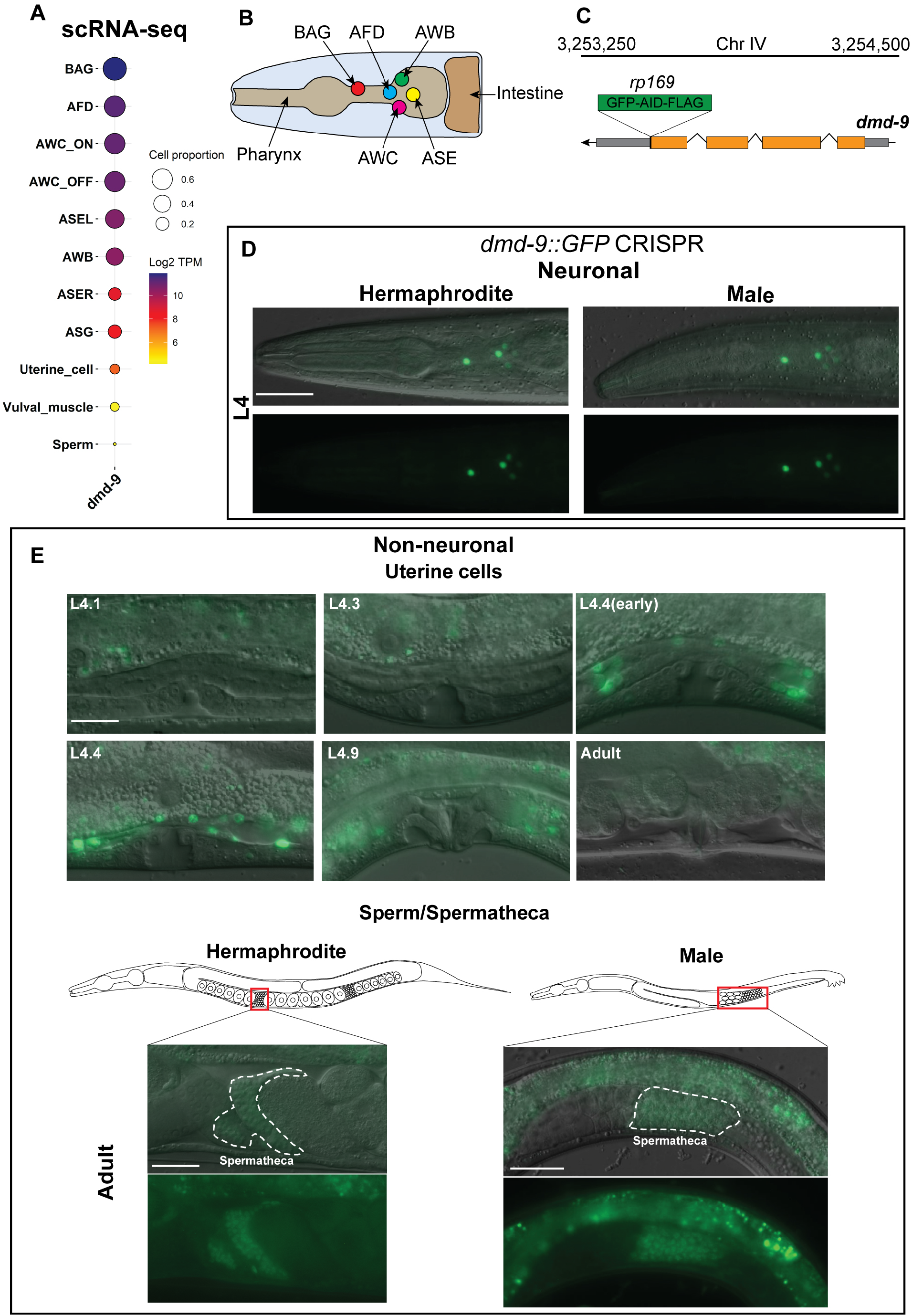
DMD-9 expression in hermaphrodites and males. **A)** Gene expression pattern of *dmd-9* in *C. elegans* obtained from (cengen.shinyapps.io/CengenApp/) (Taylor *et al*. 2021). TPM = transcript per million. **B)** Schematic of neurons that highly express *dmd-9* in the *C. elegans* head. View = left side. **C)** GFP-AID-FLAG coding sequence was endogenously inserted after the last exon of *dmd-9* gene (*rp169*) using CRISPR-Cas9 technology. **D)** Expression of *dmd-9::GFP(rp169)* in the head of hermaphrodites and males at L4 stage. Scale bar = 30 *μ*m. Anterior to the left. **E)** Expression of *dmd-9::GFP(rp169)* in non-neuronal uterine cells and spermatheca. Uterine cells exhibit transient GFP expression through the L4 stage with the highest level at L4.4 sub-stage. Scale bar = 10 *μ*m.

We confirmed the identity of DMD-9::GFP expressing head neurons by co-localization with neuron-specific reporters (Fig. 2A-B). The following reporters were used in this study: *Pets-5::mCherry* (BAGs); *Pflp-6::RFP* (AFDs); *Podr-1::RFP* (AWBs); and *Pceh-36::RFP* (AWC^on^, AWC^off^, ASEL, and ASER) (Lanjuin *et al*. 2003; Kim and Li 2004; Koga and Ohshima 2004). In addition, we found that in a small proportion of L4 animals DMD-9::GFP shows asymmetric expression in the ASEs and AWCs neurons (Fig. S1). Taken together, our results confirm the expression of DMD-9 in the neuronal cells identified by single cell transcriptomic analysis (Taylor *et al*. 2021). We did not detect sexually dimorphic expression for DMD-9::GFP within the nervous system, and observed expression in the uterine cells and spermatheca of hermaphrodites, and spermatheca in males.

**Fig 2.**
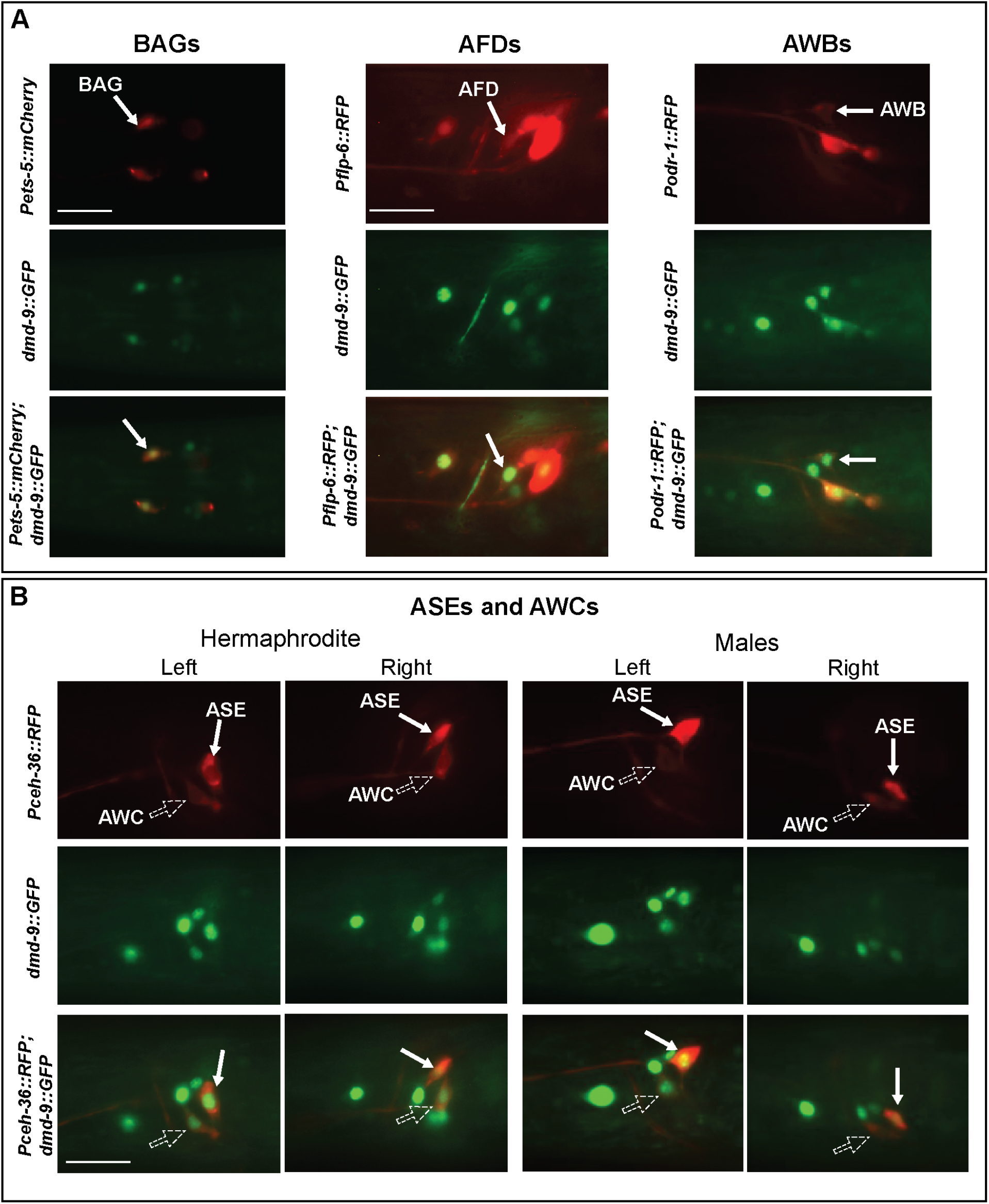
Co-localization of *dmd-9::GFP* and neuron specific reporters. **A)** Co-localization of *dmd-9::GFP* with neuron-specific markers: BAG (*Pets-5::mCherry*), AFD (*Pflp-6::RFP*) and AWB (*Podr-1::RFP*). Scale bar = 20 *μ*m. **B)** Co-localization of *dmd-9::GFP* with neuron specific marker: AWC and ASE (*Pceh-36::RFP*). Scale bar = 20 *μ*m. Strains are: *Pets-5::mCherry(RJP3088); dmd-9::GFP(rp169); Pflp-6::RFP(otIs494); Podr-1::RFP(oyIs44); Pceh-36::RFP(otIs151)*

### DMD-9 cell-autonomously regulates *flp-19* expression in the BAGs

To identify potential DMD-9 functions we obtained two deletion alleles: *dmd-9(tm4583)* and *dmd-9(ok1438)* (Fig. 3A) and crossed these strains into 15 reporters that are expressed in DMD-9 expressing neurons (Table S3). We particularly focused on the BAGs by analysing six reporters for the following genes: *flp-13, flp-17, flp-19, gcy-9, gcy-31* and *gcy-33*. To identify sexual-dimorphic gene regulation, we analysed the reporters in both hermaphrodites and males at L4 and adult stages (Table S3). Our results show that of all the reporters examined only *Pflp-19::GFP* is lost in the BAG neurons in *dmd-9* mutant animals (Fig. 3B). We identified no defects in other *flp-19* expressing neurons (Fig. 3B). We also observed dim *Pgcy-9::GFP* expression in <5% of *dmd-9* mutant animals (Table S3).

**Fig 3.**
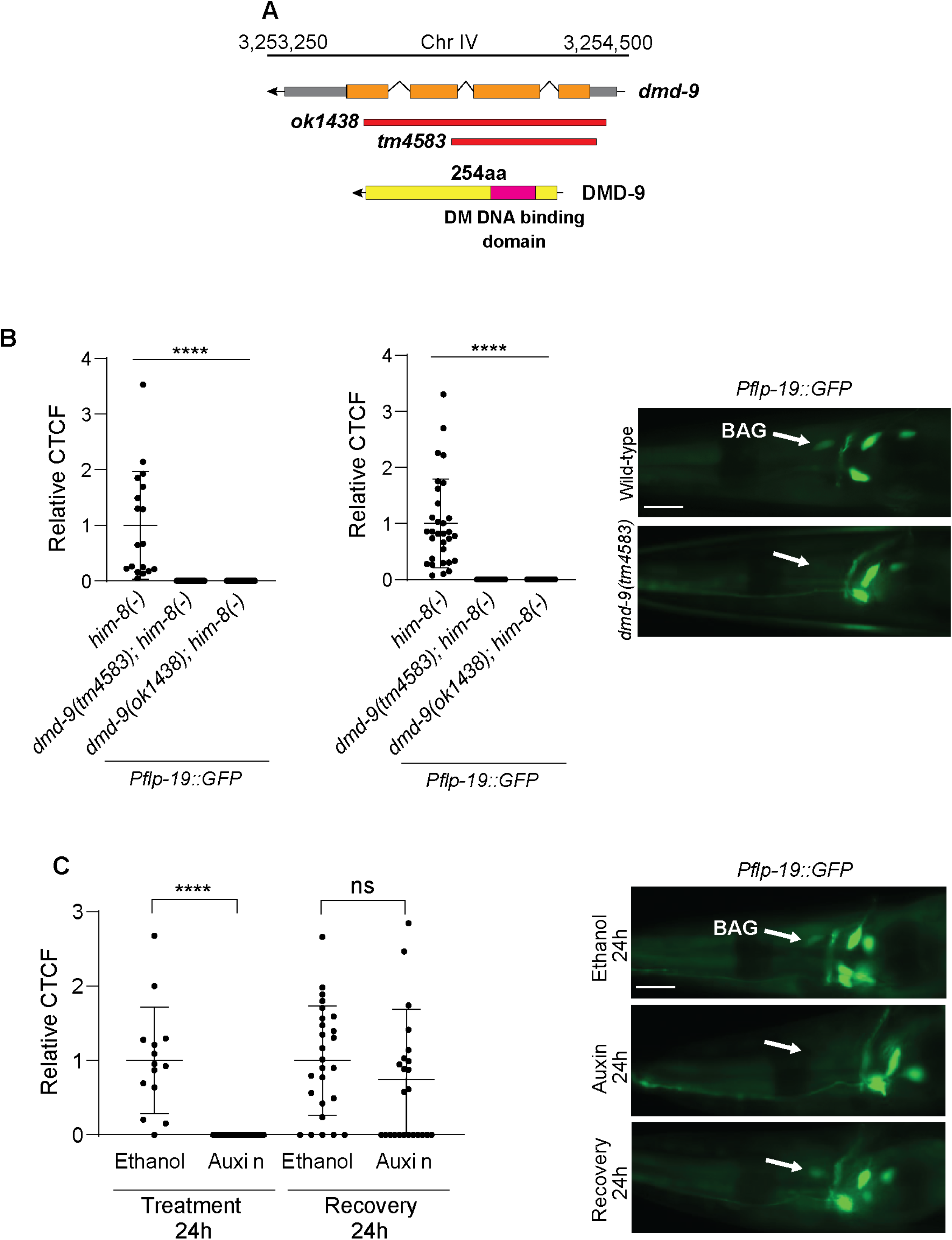
*flp-19* regulation by DMD-9. **A)** Genomic structure of the *dmd-9* gene, the available deletion mutants and protein structure with identified Doublesex/MAB-3 (DM) DNA binding domain (pink). **B)** *flp-19::GFP* reporter is not expressed in either *dmd-9* deletion allele. Scale bar = 20 *μ*m. **C)** Depletion of DMD-9 from the BAGs causes BAG-specific loss of *Pflp-19::GFP* expression. *Pflp-19::GFP* expression recovered after 24h of auxin removal. Scale bar = 20*μ*m *him-8(-)* is *him-8(e1489)*. Experiments were performed in 3 biological replications. *n* > 15, *\*\*\*\** = *P-value* ≤ 0.0001, ns = not significant. The statistical significance was calculated using one-way ANOVA with Dunnet’s correction and unpaired t-test. The error bars show SD.

To determine how DMD-9 regulates *flp-19*, we applied the AID system to deplete DMD-9 specifically in the BAG neurons. We crossed the *dmd-9::GFP::AID* strain into the *Pgcy-9::TIR1* strain that expresses the TIR1 protein only in the BAGs (Pocock *et al*. 2022). We confirmed that DMD-9::GFP is only depleted from BAG neurons and not from other neurons (Fig. S2). We then introduced the *dmd-9::GFP::AID; Pgcy-9::TIR1* strain into the *Pflp-19::GFP* reporter and performed auxin depletion. We found that after 24h of auxin-induced DMD-9::GFP depletion, expression of *Pflp-19::GFP* was undetectable in the BAG neurons (Fig. S2B). To reveal if the continual presence of DMD-9::GFP is required for *flp-19* expression we performed an auxin recovery experiment by applying a low concentration of auxin (0.1mM). The lower auxin concentration also depleted DMD-9::GFP and within 2-3 h after auxin removal DMD-9::GFP was restored (Fig. S2C). We repeated this experiment for *Pflp-19::GFP* expression. After 24h treatment with auxin (0.1mM) from the L4 stage, *Pflp-19::GFP* expression significantly reduced in the BAGs (Fig. 3C) and *Pflp-19::GFP* expression recovered after 24h following auxin removal (Fig. 3C). These results show that DMD-9 is cell-autonomously required for continuous *Pflp-19::GFP* expression in the BAGs. However, loss of DMD-9 does not adversely affect the expression of any other neuronal reporter we analyzed.

### DMD-9 has no detectable behavioral function

We investigated the function of DMD-9 in *C. elegans* behavior by focusing on the behaviors controlled by DMD-9 expressing neurons. For the majority of behaviors, we performed assays in both hermaphrodites and males to identify potential sexually-dimorphic regulatory mechanisms controlled by DMD-9.

#### Exploration and mate searching behavior

Exploration is controlled by multiple neurons, including the BAGs (Kniazeva *et al*. 2015; Juozaityte *et al*. 2017). We measured hermaphrodite exploration by tracking animal movement over a 16 h period. In contrast, we assayed male exploration over 2 and 3 h periods. In addition, we assayed male mate searching (food-leaving) behavior (Lipton *et al*. 2004; Barrios *et al*. 2008; Barrios *et al*. 2012). Our analysis of both hermaphrodites and males show that DMD-9 is not required for exploration or food-leaving behaviors (Fig. 4A and Fig. S3A).

**Fig 4.**
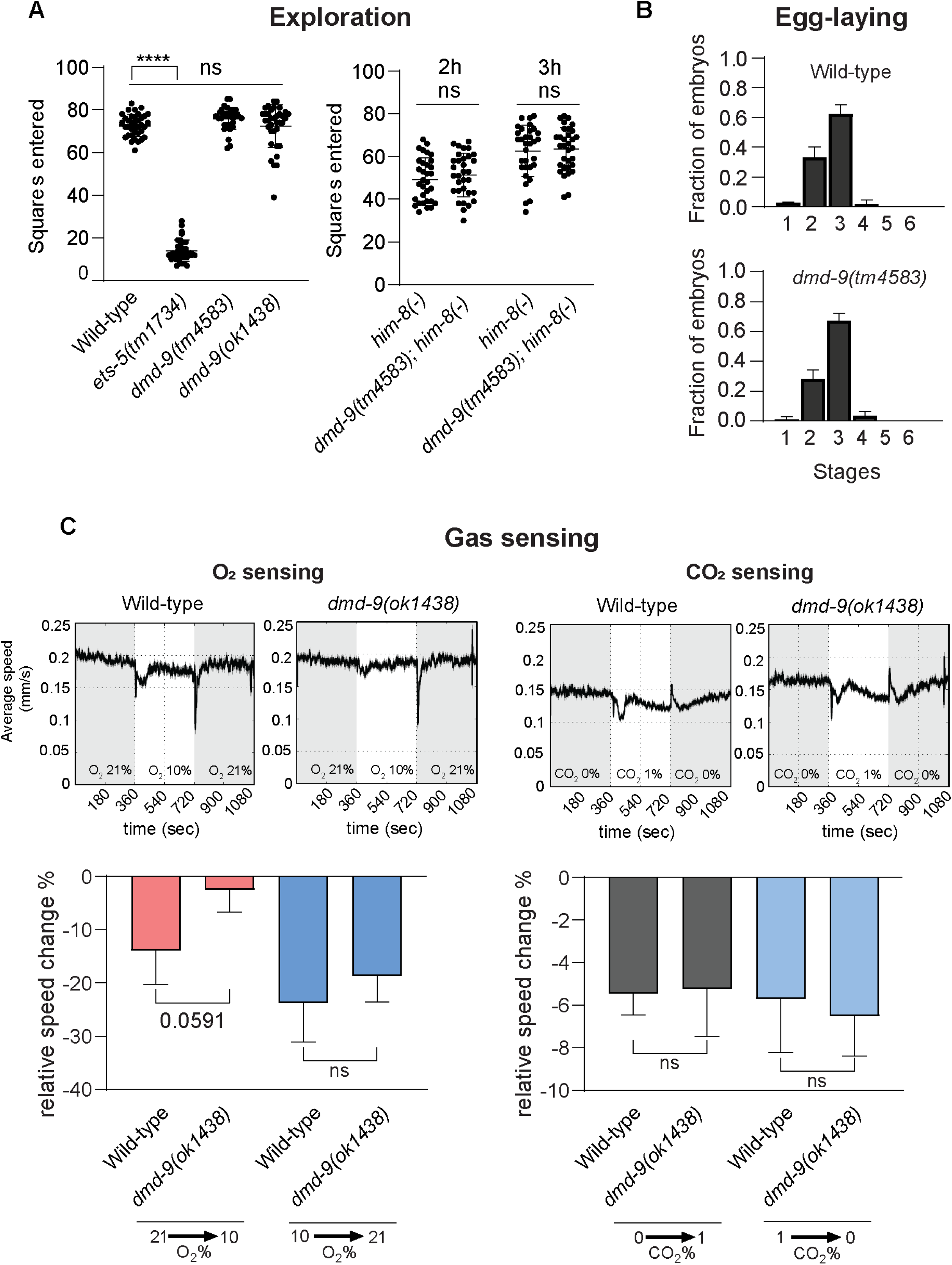
Role of DMD-9 in the BAG-mediated behaviors. **A)** Exploration behavior of L4 hermaphrodites over 16 h and males for 2 and 3 h. *ets-5(tm1734)* mutant animals used as an exploration-defective control in hermaphrodites. **B)** Egg-laying behavior. The X axis show the stage of eggs according to (Ringstad and Horvitz 2008). There was no statistical significance in egg-laying behavior between wild-type and *dmd-9(tm4583)* mutant animals. **C)** Gas sensing behavior. O_2_ and CO_2_ sensing in late L4/young adult hermaphrodites. *him-8(-)* = *him-8(e1489)*. Experiments were performed in 3 biological replications. For **A** *n* >30, **B** *n* >100, **C** *n* >400. *\*\*\*\** = *P-value* ≤ 0.0001, ns = not significant. For **A** one-way ANOVA with Dunnet’s correction and unpaired t-test, For **B** Wilcoxon Mann-Whitney rank-sum test, and for **C** unpaired t-test was used to calculate statistical significance. The error bars show SD.

#### Egg-laying

Egg-laying is regulated by multiple neurons, including the BAGs, and tissues such as the vulva and uterus (Trent *et al*. 1983; Schafer 2005; Ringstad and Horvitz 2008). As DMD-9 is expressed in the BAGs and uterine cells we analysed egg-lying in *dmd-9(tm4583)* mutant animals. Our results did not show any difference in egg-laying behavior between wild-type and *dmd-9(tm4583)* animals (Fig. 4B).

#### Gas sensing behavior

BAG neuron function is required for sensing O_2_ and CO_2_, therefore, we investigated the function of DMD-9 in sensing these gases (Hallem and Sternberg 2008; Zimmer *et al*. 2009). The experiment was performed by analyzing animal motility in response to changes in gas concentration within a sealed chamber (Romanos *et al*. 2015). Animals were exposed to switching between air (79% N_2_ and 21% O_2_) and 10% O_2_ or 1% CO_2_. The results show that *dmd-9(ok1438)* mutants did not show significant defects in O_2_ or CO_2_ sensing (Fig. 4C).

#### Chemosensory behaviors

Chemosensation is regulated by multiple neurons including the AWBs, AWCs, and ASEs (Bargmann and Horvitz 1991; Bargmann *et al*. 1993; Troemel *et al*. 1997). Since all of these neurons express DMD-9, we investigated the role of DMD-9 in regulating chemotaxis in both hermaphrodites and males. We applied high and low concentrations of each chemical and studied animal responses (Fig. S3B shows responses to high concentrations of chemicals - low concentration not shown as results were similar). We analyzed chemotaxis against the repellent 2-nonanone that is sensed by the AWBs (Troemel *et al*. 1997) (Fig. S3B). We found that *dmd-9(tm4583)* animals responded as wild-type to 2-nonanone (1:10 or 1:100 dilutions). Interestingly, wild-type males were not repelled by 2-nonanone at either concentration, suggesting sex-specific repulsion to this odor (Fig. S3B). Tanner and colleagues identified that males were repelled from undiluted 2-nonanone similar to hermaphrodites. However, males showed a delayed food-leaving behavior in the presence of 2-nonanone and food, indicating an impact of 2-nonanone on decision making (Tanner *et al*. 2022). To assess ASE neuron function, we examined chemotaxis towards 1M and 0.25M of NaCl. Our results also did not show any defects in *dmd-9(tm4583)* mutant animals (Fig. S3B). For AWC neuron function, we studied isoamyl alcohol (1:10 and 1:100) and benzaldehyde (1:1000 and 1:10000) attraction. *dmd-9(tm4583)* mutant animals exhibited wild-type attraction to these odors (Fig. S3B). Taken together, our analysis shows that DMD-9 loss does not impact behaviors regulated by neurons within which it is expressed.

### DMD-9 regulation by neuron fate determination factors

We investigated whether neuronal fate determination factors regulate the expression of DMD-9 in head neurons. We crossed *dmd-9::GFP(rp169)* into 7 fate determination TF mutants: *ets-5(tm1734/tm866)* and *egl-13(ku194)* for the BAGs; *che-1(ot866)* for the ASEs; *ttx-1(p767)* for the AFDs; *lim-4(yz12)* for the AWBs; *mls-2(tm252)* and *ceh-37(ok642)* for the AWCs (Sagasti *et al*. 1999; Satterlee *et al*. 2001; Lanjuin *et al*. 2003; Uchida *et al*. 2003; Kim *et al*. 2010; Guillermin *et al*. 2011; Gramstrup Petersen *et al*. 2013). Moreover, we investigated whether DMD-9 regulates the expression of these TFs (Table S4) and whether DMD-9 autoregulates by analysing the expression of two transcriptional GFP reporters driven by *dmd-9* promoters.

We found that four fate determining TFs regulate DMD-9 expression in different neurons. In the BAG neurons, the ETS-5 and EGL-13 TFs are required for DMD-9::GFP expression (Fig. 5A). Fosmid rescue restored DMD-9::GFP expression in the respective mutants (Fig. 5A). Loss of CHE-1 also caused loss of DMD-9::GFP expression in the ASE neurons in both hermaphrodites and males (Fig. 5B). In the AFD neurons, TTX-1 is partially required for DMD-9::GFP expression, with >50% of AFD neurons exhibiting dim expression at the L4 stage of both sexes; a defect which is exacerbated as animals become adult (Fig. 5C and Fig. S4). In contrast, we did not observe any reduction in DMD-9::GFP expression with loss of MLS-2, CEH-37 and LIM-4 – TFs required for AWBs and AWCs neuron specification (Table S5). Finally, we found that DMD-9 does not regulate the expression of any of the fate determination factors studied, and there is no evidence of DMD-9 autoregulation (Table S4). Taken together, our results show that DMD-9 expression is regulated by the ETS-5 and EGL-13 TFs in the BAGs, TTX-1 in the AFDs, and CHE-1 in the ASEs. DMD-9 does not regulate the fate of regulatory TFs or show autoregulation.

**Fig 5.**
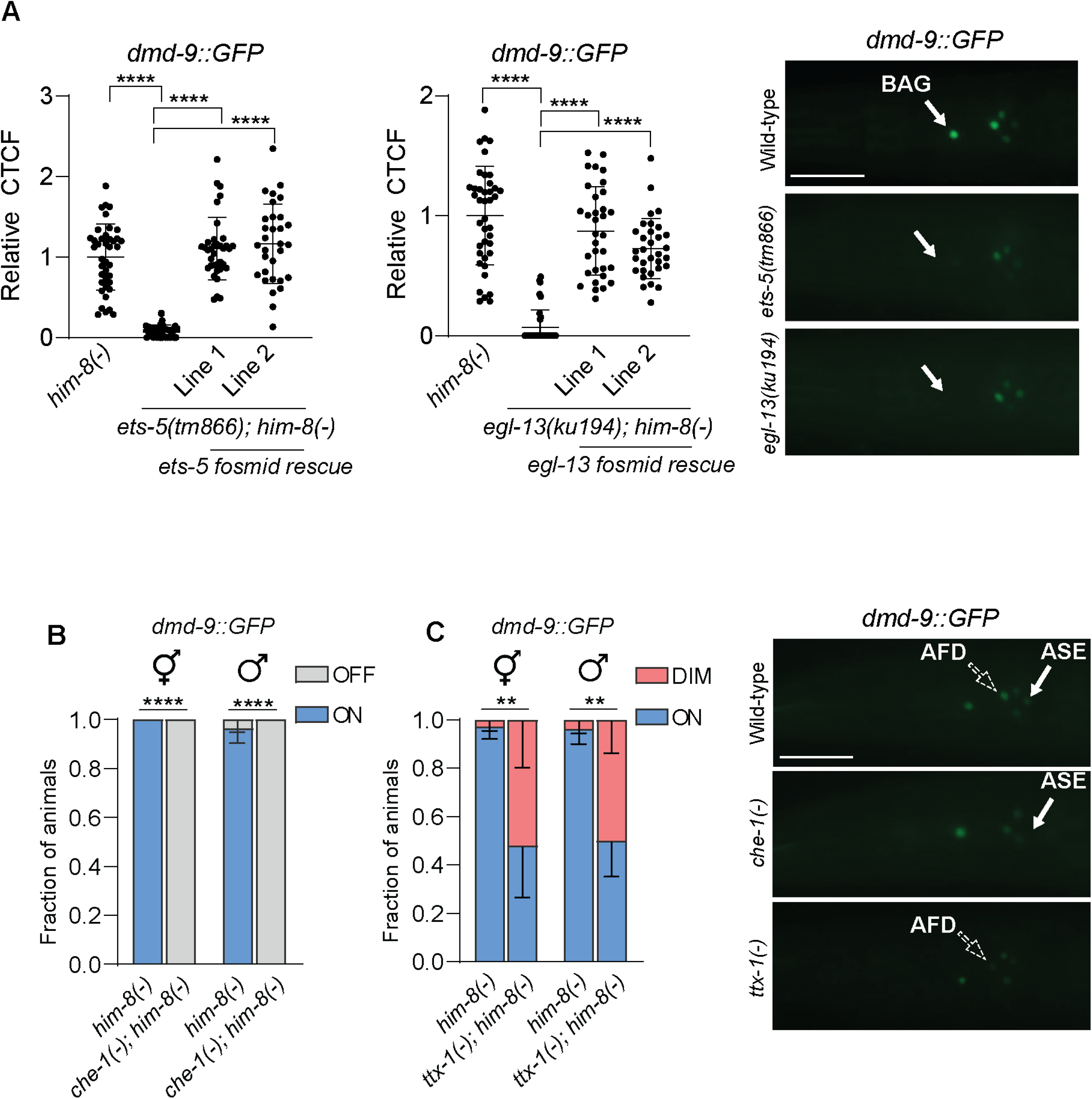
DMD-9 regulation by fate determination TFs. **A)** *dmd-9::GFP(rp169)* regulation by ETS-5 and EGL-13 TFs in L4 stage hermaphrodites. Comparisons between samples are based the relative expression of GFP to wild-type. Fosmid rescue lines significantly increased the expression of *dmd-9::GFP(rp169)* in mutant animals. Scale bar = 30 *μ*m. **B** and **C)** regulation of *dmd-9::GFP(rp169)* by CHE-1 and TTX-1 TFs. Stacked bar plots show comparisons between the wild-type and mutants at the L4 stage in both hermaphrodites and males. Scale bar = 30 *μ*m. *him-8(-)* is *him-8(e1489)*; *che-1(-)* is *che-1(ot866)*; *ttx-1(-)* is *ttx-1 (p767)*. CTCF = Corrected Total Cell Fluorescence. Experiments were performed in 2 or 3 biological replications. For **A** the significance of the results was obtained by one-way ANOVA with Dunnet’s correction for the **A** and two-way ANOVA analyses with comparing simple effect within columns for the **B** and **C**. *n* > 30, *\*\*\*\** = *P-value* ≤ 0.0001, *\*\** = *P-value* ≤ 0.01, ns = not significant. The error bars show SD.

### *cis*-regulatory analysis of the *dmd-9* promoter

We have shown that DMD-9 is regulated by multiple neuron-specific fate determining TFs (Fig. 5). Based on these observations, we examined which regulatory elements within the *dmd-9* promoter direct neuron-specific expression. To investigate this, we used a fluorescent reporter driven by a 4.5kb *dmd-9* promoter that drives GFP in the BAGs, neurons posterior to the nerve ring, and uterine cells (Fig. 6). We truncated this template reporter by PCR to generate 3kb, 2kb, 1kb, 500bp, 450bp, 375bp, 250bp, and 100bp versions of the promoter to drive GFP (Fig. 6). Our results show that the 500bp promoter (*prom5*) is sufficient to drive GFP in the same neurons as the full promoter (4.5kb). However, the *dmd-9prom8::GFP* (250bp promoter) was no longer expressed in the BAGs, but maintained expression in neurons posterior to the nerve ring. The *dmd-9prom9::GFP* (100bp promoter) is not expressed in any neurons (Fig. 6). We focused on the regulatory elements in the *dmd-9* promoter that control GFP expression in the BAGs by further dissecting the 500-250bp region into *dmd-9prom6::GFP* (450bp) and *dmd-9prom7::GFP* (376bp). We found that *dmd-9prom7::GFP* drives GFP in the BAG neurons at reduced intensity compared to *dmd-9prom6::GFP*. We constructed a GFP reporter (*dmd-9prom10::GFP*) by cloning 500-250bp segments of the *dmd-9* promoter into the pPD95.75 GFP vector and found that it drives robust GFP expression in the BAG neurons (Fig. 6). Analysis of this sequence in JASPAR (jaspar.genereg.net) identified two potential regulatory elements: two Ets sites and one Otx2 site (Fig. 6). Site-directed mutagenesis of the Ets sites showed no overt effect on the GFP expression in the BAG neurons (Fig. 6). In addition, the *dmd-9prom10::GFP* was expressed in *ets-5(tm1734)* mutant animals (Table S6). It is therefore likely that ETS-5 regulates *dmd-9* expression in the BAGs indirectly or in combination with other factors (Fig. 5A). We also mutagenized the Otx2 site alone or in combination with the Ets mutated sites. We found that GFP expression of *dmd-9prom10::GFP* is completely lost in BAG neurons following mutagenesis of the Otx2 site. Potential TFs that may bind to Otx2 elements include the OTX TFs CEH-36, CEH-37 and TTX-1, however, these TFs are not expressed in the BAG neurons (Lanjuin *et al*. 2003). Therefore, we searched for other homeobox TFs expressed in the BAGs that potentially regulate DMD-9 through the Otx2 element. We identified the CEH-23 and CEH-54 homeobox TFs as candidates and introduced the *dmd-9prom10::GFP* construct into *ceh-23(ms23)* and *ceh-54(tm242)* mutants. We found no change in GFP expression in the BAGs compared to wild-type animals (Table S6). Taken together, *cis-*regulatory analysis of the *dmd-9* promoter identified a 500bp upstream region that is sufficient for head neuron expression. We found that at least one Otx2 element is required to drive BAG expression, however, the identity of the factors regulating this site are unknown.

**Fig 6.**
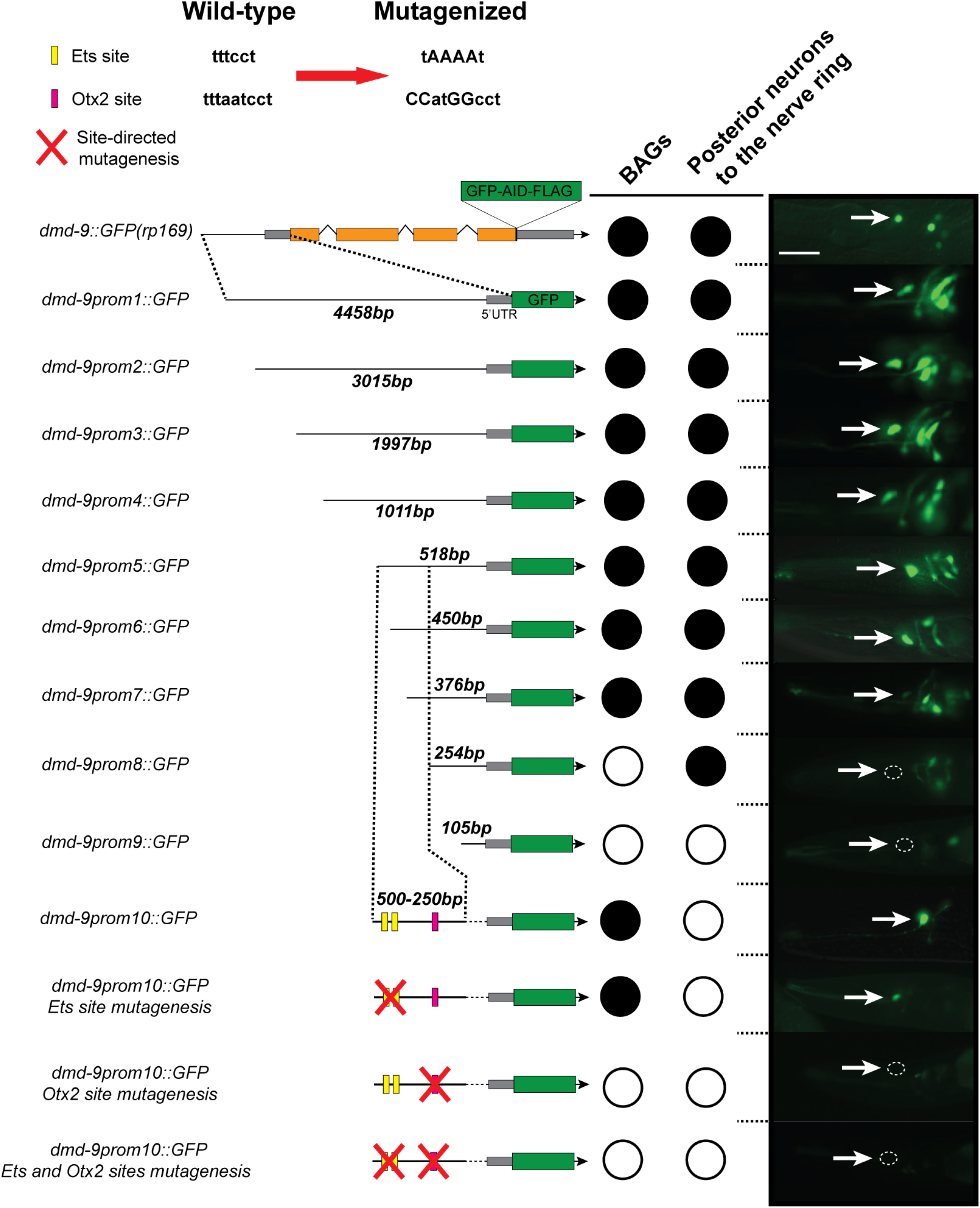
*dmd-9* promoter dissection analysis. We used promoter deletion and mutagenesis to identify regions and elements that regulate *dmd-9* expression in the BAGs and the posterior neurons to the nerve ring. For each constructed reporter, 3 independent lines and more than 20 animals were analyzed. White arrows show BAG neurons location. Scale bar = 20 *μ*m.

## Discussion

The *C. elegans* DMD family encompasses 11 TFs, some of which control sexually-dimorphic characteristics (Shen and Hodgkin 1988; Yi *et al*. 2000; Lints and Emmons 2002; Oren-Suissa *et al*. 2016; Serrano-Saiz *et al*. 2017; Bayer *et al*. 2020; Durbeck *et al*. 2021). Here we analyzed another member of this family, DMD-9. According to single-cell transcriptomic data (Taylor *et al*. 2021) *dmd-9* is expressed in multiple sensory neurons and non-neuronal tissues. We studied the expression of DMD-9 by generating a CRISPR-tagged GFP reporter and confirmed the *dmd-9* expression results from single cell RNA sequencing (Taylor *et al*. 2021). We did not identify sexual-dimorphic expression of *dmd-9*, however, expression in the hermaphrodite specific uterine cells and spermatheca was detected. DMD-9 may have a specific function in the development of the uterus as expression occurs during a sub-stage of L4 (Fig. 1E). We did not observe any obvious defects in uterine morphology or egg-laying behavior that is partially regulated by uterine cells (Schafer 2005). Another DMD family member, MAB-23 is also expressed in cells related to egg-laying, including the uterus (Lints and Emmons 2002). Therefore, multiple DMD TFs may act redundantly to control the fate and/or function of uterine cells – a concept requiring future study. DMD-9 is also expressed in the spermatheca of hermaphrodites and males. However, there are no obvious defects in the mating process in *dmd-9* mutant males.

According to our analysis, DMD-9 is not broadly involved in nervous system fate determination. We found that DMD-9 cell-autonomously regulates *flp-19*, which encodes a FMRF-like peptide that is expressed in multiple neurons including the BAGs and AWAs. However, the function of FLP-19 is poorly understood. Elevated FLP-19 expression in the context of an *arcp-1* mutant enhances behavioral responses to CO_2_ (Beets *et al*. 2020). FLP-19 is also required for exploration behavior (Juozaityte *et al*. 2017). In our study, *dmd-9* mutants behave similar to wild-type in both CO_2_ sensing and exploration. Therefore, loss of FLP-19 in the BAG neurons in *dmd-9* mutant animals may cause defects that are yet to be identified.

Analyzing behaviors that could potentially be mediated by *dmd-9*-expressing neurons did not reveal any defects compared to wild-type animals. Studies of other members of the DMD family have shown a wide range of phenotypes including sex-specific behavior and morphological defects (Shen and Hodgkin 1988; Yi *et al*. 2000; Oren-Suissa *et al*. 2016). Sex-specific expression patterns of DMD-3, DMD-4, DMD-5 and DMD-11 regulate sex-specific synaptogenesis in these neurons (Oren-Suissa *et al*. 2016; Serrano-Saiz *et al*. 2017; Bayer *et al*. 2020). DMD-5 and DMD-11 also regulate AVG-mediated male mating behavior (Oren-Suissa *et al*. 2016). Regarding sex shared behavior, DMD-10 regulates ASH-mediated responses to nose touch and high osmolarity (Durbeck *et al*. 2021). These findings suggest that DMD TFs, (except MAB-3 and MAB-23 which show defects in the morphology of the male animals (Shen and Hodgkin 1988; Yi *et al*. 2000; Lints and Emmons 2002)) do not show severe defects in sex-shared phenotypes, however, more studies are required to identify these associations. The lack of behavioral deficits in *dmd-9* mutant animals suggest subtle phenotypes that cannot be identified by current conventional experiments, or alternatively functional redundancy with other genes which compensate for DMD-9 loss. DMD-4, which is expressed in several head neurons and loss of which is lethal (Bayer *et al*. 2020) and DMD-6 which shows pan-neuronal expression (cengen.shinyapps.io/CengenApp/)(Taylor *et al*. 2021) are candidate redundantly acting factors. In addition, the role of DMD-9 in AFD-mediated thermosensory behavior needs to be examined.

DMD-9 is expressed in specific head neurons and is independently regulated by fate determining TFs in each neuron. We found that in the BAGs, AFDs, and AWCs, *dmd-9* is regulated by cell-specific terminal regulatory TFs. In the BAGs, ETS-5 and EGL-13 regulate neuronal fate by controlling the expression of multiple genes such as *flp-17, gcy-9, gcy-31, gcy-33* (Guillermin et al. 2011; Gramstrup Petersen et al. 2013). TTX-1 regulates gene batteries of the AFD neurons (Satterlee *et al*. 2001) and the fate and functionality of ASE neurons is regulated by CHE-1 (Uchida *et al*. 2003). Noticeably, *dmd-9* regulation by these fate determining factors is neuron-specific, suggesting cell-specific regulatory mechanisms or the presence of TF-specific *cis-*regulatory elements. In the BAG neurons, we found that at least one Otx2 element within 400bp upstream of the *dmd-9* start codon drives GFP expression. In addition, elements within 250bp are required for GFP expression in the neurons posterior to the nerve ring. Otx2 elements may be regulated by CEH-36, CEH-37 and TTX-1 Otx homeodomain TFs (Lanjuin *et al*. 2003), however they are not expressed in the BAG neurons. Searching for other potential regulators through the Otx2 site, we analyzed the CEH-23 and CEH-54 homeodomain TFs that are expressed in the BAG neurons. However, loss of these TFs did not affect DMD-9 expression in the BAG neurons. ETS-5 is required for *dmd-9* expression in the BAGs but shows no effect on the transcriptional reporter carrying Ets and Otx2 elements. Therefore, ETS-5 and EGL-13 likely regulate DMD-9 expression indirectly through other unknown regulatory factors in the BAG neurons.

## Conclusion

In this study, we characterized the DMD-9 TF in both hermaphrodite and males. We found that FLP-19 expression in the BAGs is cell-autonomously regulated by DMD-9. We also identified the *cis*-regulatory elements and *trans*-acting factors required for DMD-9 expression in specific sensory neurons.

## Acknowledgements

We thank members of the Pocock laboratory for comments on the manuscript. Some strains used in this study were provided by the *Caenorhabditis* Genetics Center, which is funded by NIH Office of Research and Infrastructure Programs (P40 OD10440) and the National BioResource Project (NBRP) Japan. This work was supported by the following funding: National Health and Medical Research Council grants GNT1105374, GNT1137645 and GNT2000766 (RP).

## Data and Materials Availability

All data are available in the main text or the supplementary materials.

## Supplementary Figure and Table Legends

**Fig S1.**Analysis of the *dmd-9::GFP(rp169)* expression in the left and right AWC and ASE neurons. Experiments were performed in 3 biological replications. n >40. We used two-way ANOVA analyses with comparing simple effect within columns to analyse significance of the results. The error bars show SD.

**Fig S2.**Depletion of DMD-9::GFP protein using the AID system.

**A)** All the animals lost DMD-9::GFP expression in the BAG neurons of animals carrying the *Pgcy-9::TIR1* transgene that were exposed to auxin.

**B)** DMD-9::GFP protein is depleted from the BAGs after 2h of auxin treatment. *Pflp-19::GFP* expression is also undetectable following 24h auxin treatment. Scale bar = 20 *μ*m.

**C)** Auxin treatment and recovery of DMD-9. Animals were exposed to auxin (0.1mM) for 2h and confirmed for depletion of *dmd-9* expression in the BAGs. A portion of animals were transferred to NGM plates and incubated for 3h and then examined for recovery of *dmd-9* expression.

White arrows show the BAG neurons. CTCF = Corrected Total Cell Fluorescence. Experiments were performed in 3 biological replications. *n* >15, *\*\*\*\** = *P*-value ≤ 0.0001, ns = not significant. The unpaired t-student test was used to obtain statistical significance. The error bars show SD.

**Fig S3.***dmd-9* behavioral analysis.

**A)** Mate searching behavior in both adult males and hermaphrodites. *pdfr-1(ok3425)* mutant used as a male leaving-deficient control.

**B)** Chemosensory behaviors regulated by AWB, AWC and ASE neurons. 2-butanone, benzaldehyde, isoamyl alcohol, and NaCl were used as attractant chemicals, while 2-nonanone was used as repellent. For these experiments, positive controls were as follows: *odr-3(n2150)* for 2-butanone, benzaldehyde, isoamyl alcohol, and 2-nonanone; *che-1(ot866)* for NaCl (not shown).

*him-8(-)* is *him-8(e1489)*. Experiments were performed in 5 biological replications. For the mate-searching behavior *n* > 60. For the chemosensory behaviors *n* >300, *\*\*\*\** = *P-value* ≤ 0.0001, ns = not significant. For **A** the Gehan-Breslow-Wilcoxon test and For **B** unpaired t-test were applied to calculate statistical significance. The error bars show SD.

**Fig S4.***dmd-9::GFP(rp169)* regulation by CHE-1 and TTX-1 TFs. Experiments were performed in 2 biological replications. *him-8(-)* is *him-8(e1489)*; *che-1(-)* is *che-1(ot866)*; *ttx-1(-)* is *ttx-1 (p767)*. The significance of the results was obtained by two-way ANOVA analyses with comparing simple effect within columns analyses. *n* >30, *\*\*\*\** = *P-value* ≤ 0.0001, *\*\*\** = *P-value* ≤ 0.001, ns = not significant. The error bars show SD.

## Supplementary Tables

**Table S1.**
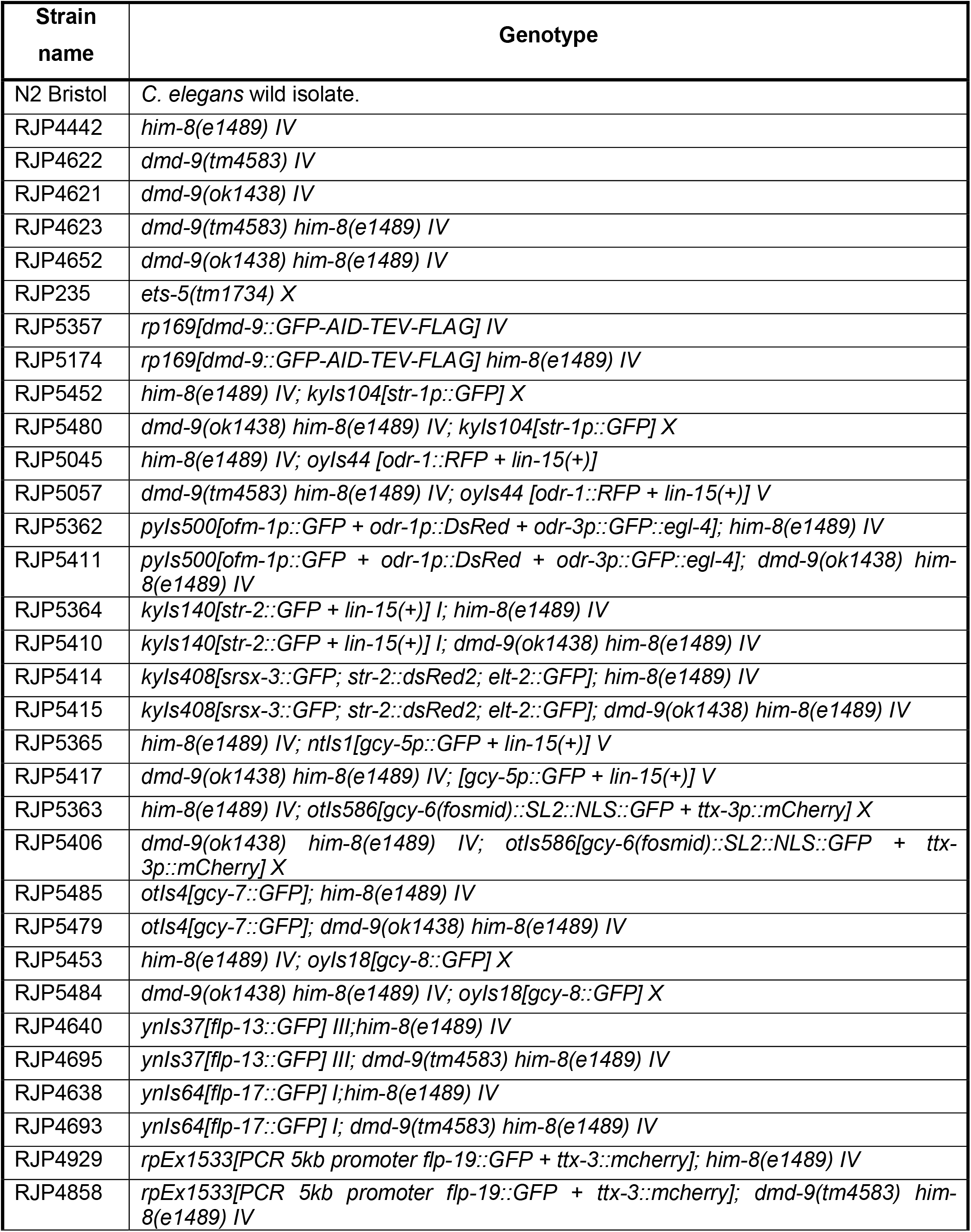
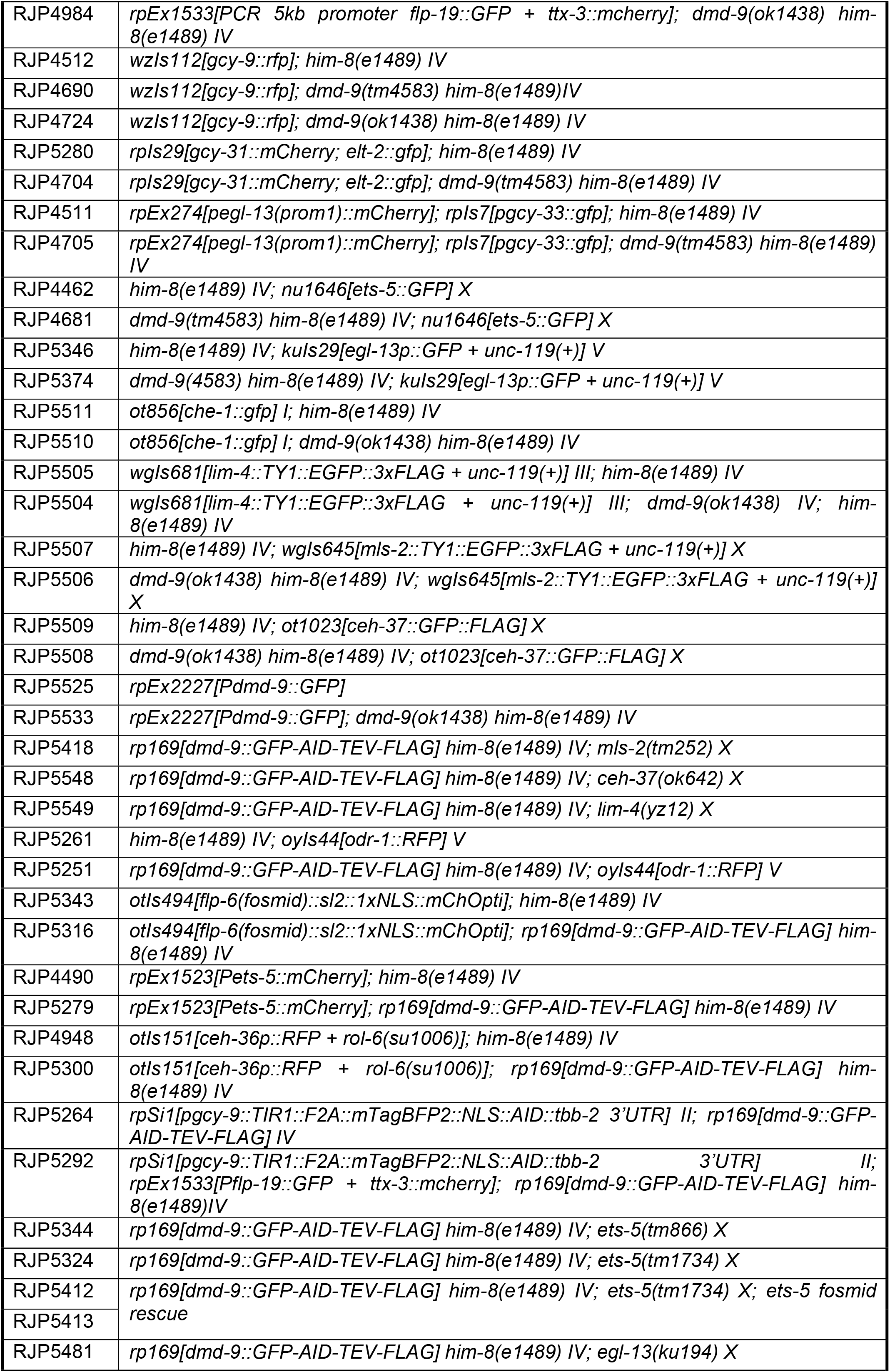
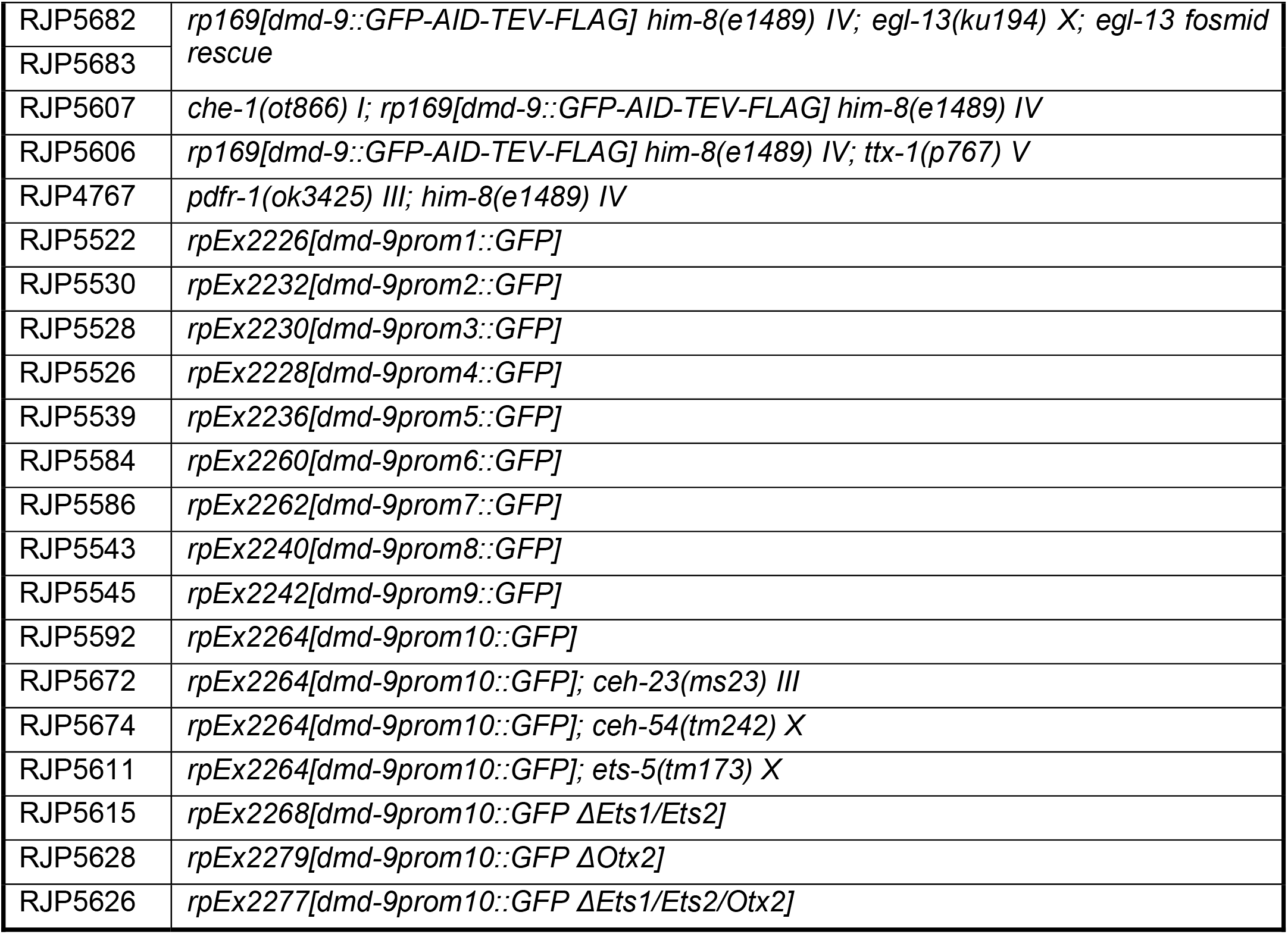
Strains used in this study.

**Table S2.**
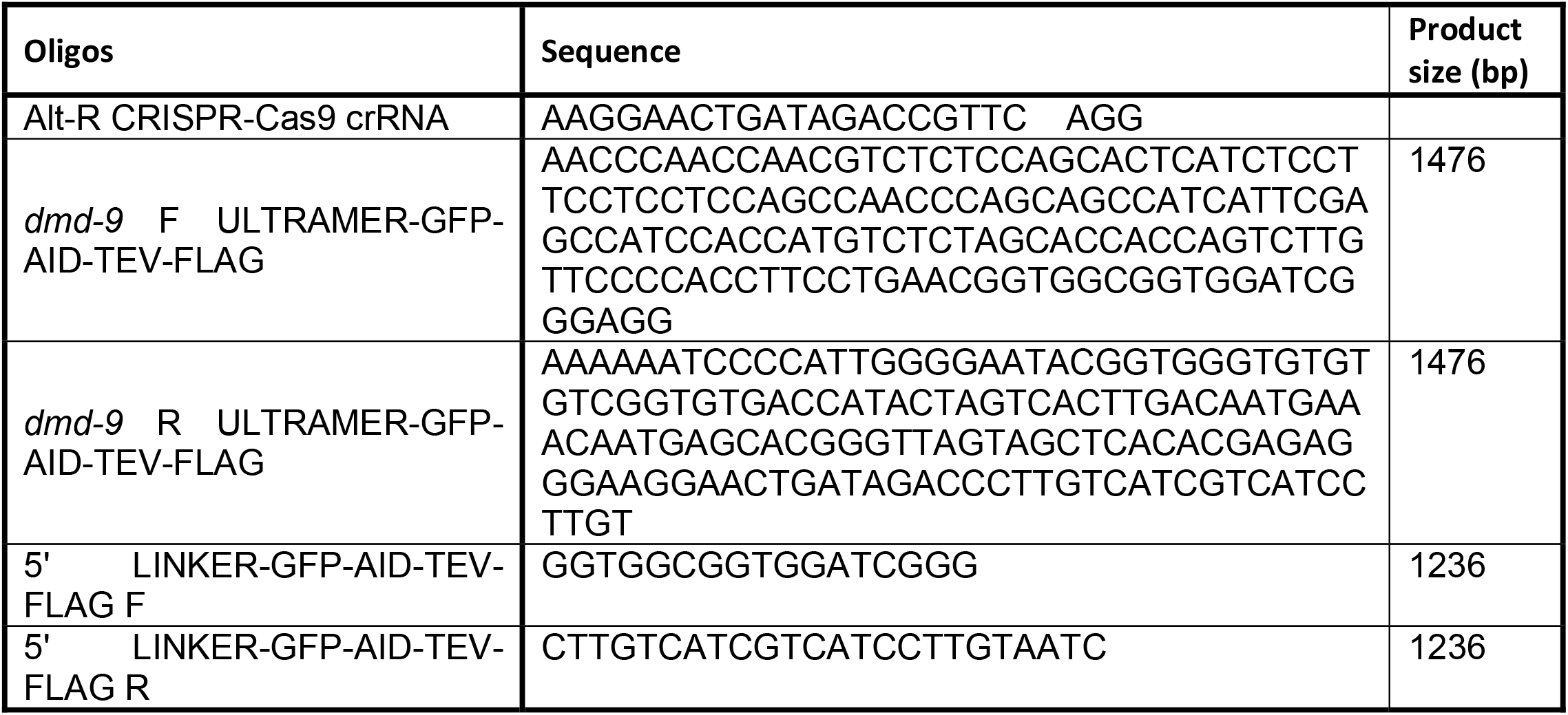
Oligos used for GFP knock-in to *dmd-9* using CRISPR-Cas9.

**Table S3.**
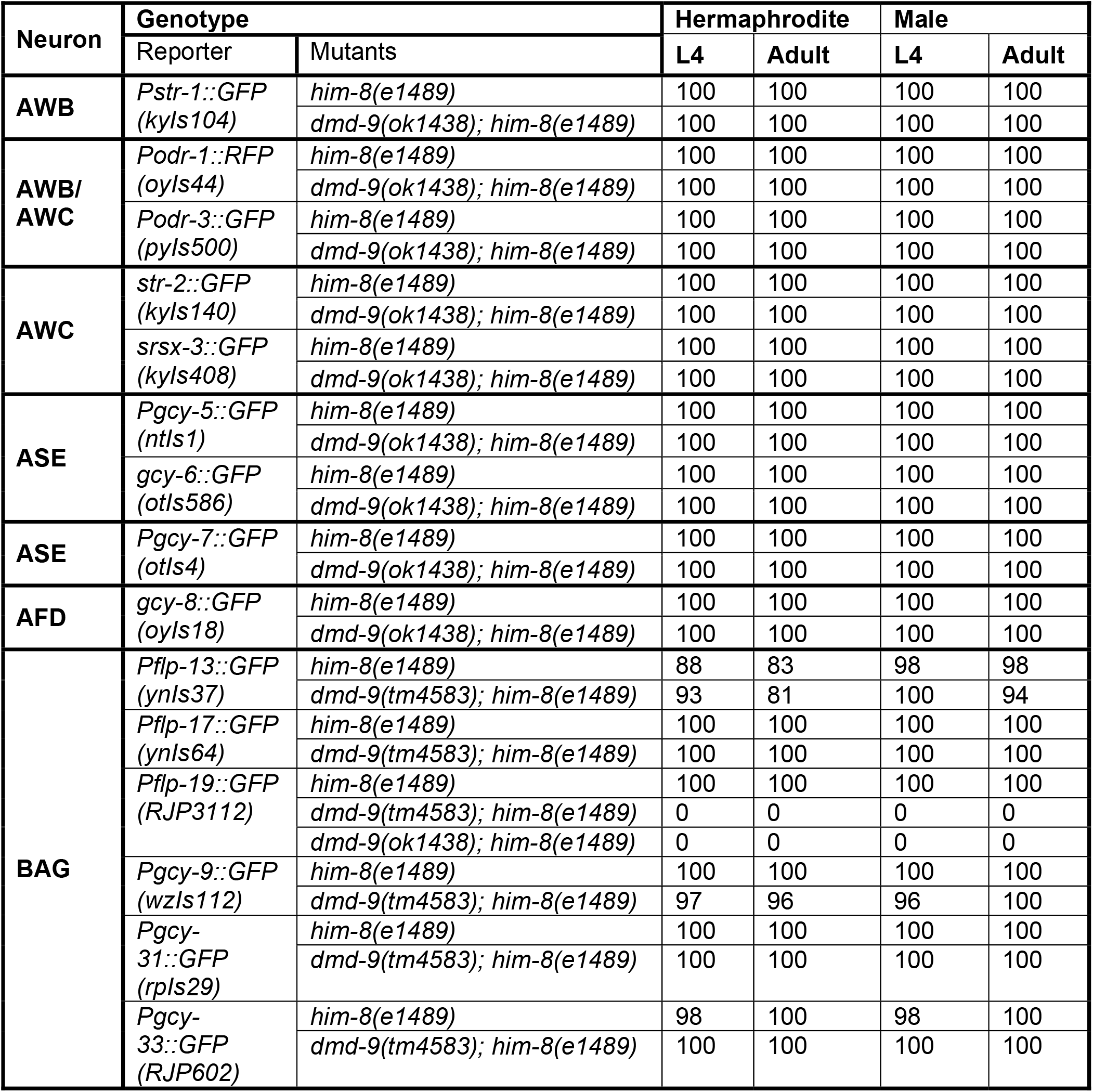
Neuron fate reporters used in this study and the impact of the DMD-9 TF on their expression in L4 and adult hermaphrodites and males. Numbers show the percentage of animals expressing the reporter.

**Table S4.** DMD-9 TF regulation of neuron fate determining TFs. Numbers show the percentage of animals expressing the reporter.

| Neuron | Genotype |  | Hermaphrodite |  | Male |  |
| --- | --- | --- | --- | --- | --- | --- |
|  | Reporter | Mutants | L4 | Adult | L4 | Adult |
| **BAG** | *ets-5::GFP*<br>(*nu1646*) | *him-8(e1489)* | 100 | 100 | 100 | 100 |
|  |  | *dmd-9(tm4583); him-8(e1489)* | 100 | 100 | 100 | 100 |
|  | *egl-13::GFP*<br>(*kuls29*) | *him-8(e1489)* | 100 | 100 | 100 | 100 |
|  |  | *dmd-9(ok1438); him-8(e1489)* | 100 | 100 | 100 | 100 |
| **ASE** | *che-1::GFP*<br>(*ot856*) | *him-8(e1489)* | 100 | 100 | 100 | 100 |
|  |  | *dmd-9(ok1438); him-8(e1489)* | 100 | 100 | 100 | 100 |
| **AWB** | *lim-4::GFP*<br>(*wgls681*) | *him-8(e1489)* | 100 | 100 | 100 | 100 |
|  |  | *dmd-9(ok1438); him-8(e1489)* | 100 | 100 | 100 | 100 |
| **AWC/ASE** | *mls-2::GFP*<br>(*wgls645*) | *him-8(e1489)* | 100 | 100 | 100 | 100 |
|  |  | *dmd-9(ok1438); him-8(e1489)* | 100 | 100 | 100 | 100 |
| **AWC/ASE/**<br>**AWB** | *ceh-37::GFP*<br>(*ot1023*) | *him-8(e1489)* | 100 | 100 | 100 | 100 |
|  |  | *dmd-9(ok1438); him-8(e1489)* | 100 | 100 | 100 | 100 |
| **BAG/AFD/**<br>**AWC/**<br>**ASE/AWB** | *rpEx2226[Pdmd-9::GFP]* | *N2* | 100 | 100 | - | - |
|  |  | *dmd-9(ok1438); him-8(e1489)* | 100 | 100 | 100 | 100 |

**Table S5.** Role of neuron fate determining TFs on *dmd-9* regulation. Numbers show the percentage of animals expressing the reporter.

| Neuron | Genotype |  | Hermaphrodite |  | Male |  |
| --- | --- | --- | --- | --- | --- | --- |
|  | Reporter | Mutants | L4 | Adult | L4 | Adult |
| AWC* | *dmd-9::GFP(rp169)* | *him-8(e1489)* | 97 | 95 | 92 | 100 |
|  |  | *mls-2(tm252); him-8(e1489)* | 95 | 100 | 97 | 100 |
|  | *dmd-9::GFP(rp169)* | *ceh-37(ok642); him-8(e1489)* | 93 | 98 | 98 | 93 |
| AWB | *dmd-9::GFP(rp169)* | *him-8(e1489)* | 100 | 100 | 100 | 100 |
|  |  | *lim-4(yz12); him-8(e1489)* | 100 | 100 | 100 | 100 |

\*: average of AWC left and right.

**Table S6.** Role of TFs on the regulation of Ets and Otx2 *cis*-regulatory elements.

| Strains | Line 1 |  |  | Line 2 |  |  |
| --- | --- | --- | --- | --- | --- | --- |
|  | ON | dim | OFF | ON | dim | OFF |
| *Wild-type* | 0.41 | 0.13 | 0.44 | 0.33 | 0.08 | 0.57 |
| *ets-5(tm1734)* | 0.33 | 0.13 | 0.52 | 0.32 | 0.12 | 0.55 |
| *ceh-23(ms23)* | 0.34 | 0.27 | 0.37 | 0.26 | 0.2 | 0.53 |
| *ceh-54(tm242)* | 0.33 | 0.11 | 0.55 | 0.31 | 0.15 | 0.53 |

